# Multi-Scale Simulations of Excited-State Energy Transfer Pathways in the C2S2-Type PSII-LHCII Supercomplex of Spinach

**DOI:** 10.1101/2025.09.14.675516

**Authors:** Jianping Guo, Fan Xu, Ruichao Mao, Lihua Bie, Luning Liu, Jun Gao

## Abstract

Photosystem II-light-harvesting complex II (PSII-LHCII) supercomplex achieves efficient excitation energy transfer (EET), yet how directional transfer emerges from its nearly flat energy landscape has remained unclear. Here, we investigate the C_2_S_2_-type spinach PSII-LHCII supercomplex containing 206 pigments by integrating microsecond-scale molecular dynamics, QM/MM site energy calculations, and generalized Förster theory. The computed site energy landscape shows overall downhill gradients from peripheral antennas to the reaction center (RC), with small variations fully explained by dynamic structural fluctuations. Pairwise transfer rates reveal a dense multi-timescale EET network, where fast intra-subunit and interface couplings provide robust channels and slower links maintain global connectivity. Key interface pigments act as energy-funneling hotspots that converge excitations toward the RC. Within the PSII core, both the D1 and D2 branches ultimately transfer energy downhill to Chl_D1_ and Chl_D2_, respectively, ensuring thermodynamic feasibility of delivery. Three major transfer pathways were identified, all falling within functional timescales (< 250 ps). Bidirectional dynamics were also observed, with rapid reverse transfers enabling excess excitation to be safely dissipated through peripheral antennas. Together, these results demonstrate that PSII maintains fully downhill energy transfer through the combined effects of structural constraints and dynamic regulation, thereby achieving both efficient energy delivery and photoprotection.

## INTRODUCTION

Photosynthesis is a fundamental biological process that converts solar energy into chemical energy, supporting life on Earth for over 3.5 billion years^1–3^. This highly efficient energy conversion relies on photosystems-sophisticated pigment-protein complexes embedded in thylakoid membranes. In oxygenic photosynthetic organisms such as higher plants and algae, photosystem I (PSI) and photosystem II (PSII) are responsible for light-driven electron transfer^4, 5^. PSII, in particular, is a multifunctional enzyme complex involved in light harvesting, water splitting, oxygen evolution, and the coupled translocation of protons and electrons^6, 7^.

The structural organization of photosynthetic machinery plays a critical role in guiding excitation energy transfer (EET), ensuring that the efficient delivery of absorbed energy to the reaction center (RC). In the photosystem, the spatial arrangement of pigments, their mutual orientations, and the surrounding protein environment collectively contribute to influencing the pathways and rates of energy transfer^8–14^. Understanding these structure-function relationships is essential for elucidating PSII mechanisms and for guiding the design of artificial photosynthetic systems, advancing solar to fuel technologies^15, 16^, and improving crop light use efficiency^17–19^. However, due to the complexity and dynamic nature of PSII, resolving its structure and dynamics under functional conditions remains challenging, and our understanding of EET is still limited, particularly regarding transient conformations and subtle structural heterogeneity that influence energy flow.

The advent of cryo-electron microscopy (cryo-EM) has enabled near-atomic resolution imaging of PSII, revealing its complex pigment-protein architecture^20^. Structural studies have identified multiple PSII-LHCII supercomplex types, such as C_2_S_2_-type, C_2_S_2_M_2_-type, and C_2_S_2_M_2_L_2_-type, differing in peripheral antenna composition across species^3, 21^. The first high-resolution spinach PSII-LHCII structure, reported in 2016, revealed a C_2_S_2_-type homodimer in which each monomer contains 25 protein subunits and 105 chlorophylls^22^. Later work resolved structures from pea and *Arabidopsis thaliana* in both C_2_S_2_-type and C_2_S_2_M_2_-type configurations, showing how additional M-LHCII subunits expand the antenna network^23–26^. In green algae, *Chlamydomonas reinhardtii* PSII-LHCII complexes have been characterized in C_2_S_2_ and C_2_S_2_M_2_L_2_ forms^27^. These studies confirm C_2_S_2_ as the core architecture of green lineage PSII-LHCII, forming the structural basis for conserved EET pathways^21^.

While static structures reveal pigment positions, subunit assembly, and cofactor binding sites, they cannot directly capture EET processes, such as excitation migration between pigments and transfer across subunit interfaces. These processes are shaped by both structural and environmental factors and require complementary approaches. Theoretical modeling and ultrafast spectroscopy have been particularly useful^28, 29^. Multiscale simulations combining molecular dynamics (MD) and quantum mechanics/molecular mechanics (QM/MM) calculations have been used to study energy transfer within isolated PSII subunits, revealing how local structure affects excitation energies and couplings^28, 30^. However, these studies often focus on individual components rather than the full complex. QM-based analyses on PSII from pea and green algae have provided organism-specific insights into pigment-protein interactions and environmental effects on EET^31, 32^. Despite these advances, comprehensive all-atom studies of PSII EET encompassing both intra- and inter-subunit dynamics are still limited, leaving aspects of its complete pigment network insufficiently understood.

The energy landscape of PSII is a key factor in understanding its function. While many photosystems have a clear downhill “funnel” to the RC^33^, PSII has a more complex, flatter profile lacking a single dominant funnel^34, 35^. How this near-flat gradient directs efficient, directional EET is unresolved^36^. Protein network topology also plays a role but is poorly understood^4^. In 2016, Kreisbeck at al. used QM/MM simulations on a 12 Å structure of the PSII-LHCII supercomplex and found that EET is not purely downhill and may encounter a barrier before the RC^37^. Low-resolution structural limitations, however, may have influenced these results. On the experimental side, two-dimensional electronic spectroscopy (2DES) has shown that PSII’s flat energy landscape supports bidirectional energy flow between core and peripheral antennas^38–40^. This bidirectionality not only facilitates efficient light harvesting under low-light conditions but also underpins the photoprotective mechanisms activated under high-light stress^41^. Despite these advances, no consensus model of PSII EET exists, and the role of its energy landscape and network topology remains debated^36, 42, 43^.

In this work, we investigate the C_2_S_2_-type spinach PSII-LHCII supercomplex^22^. The arrangement of its pigments and protein subunits is shown in Fig. 1. This complex is made up of two monomers, with each monomer containing a core complex. The core complex consists of four large intrinsic subunits: D1, D2, CP43, and CP47. Adjacent to CP43 within the core complex, one LHCII trimer and one CP26 subunit are located. On the opposite side, a CP29 subunit interacts with CP47. The complex contains 206 pigments. Among these, 156 are chlorophyll a distributed across multiple subunits, and 50 are chlorophyll b, found only in LHCII, CP29, and CP26. Previously, based on this structure, our group constructed a million-atom scale model of this supercomplex embedded in the solvated membrane and performed microsecond-scale molecular dynamics simulations to study the protein interactions and assembly process of the sizeable PSII-LHCII supercomplex^44^.

**Fig. 1.**
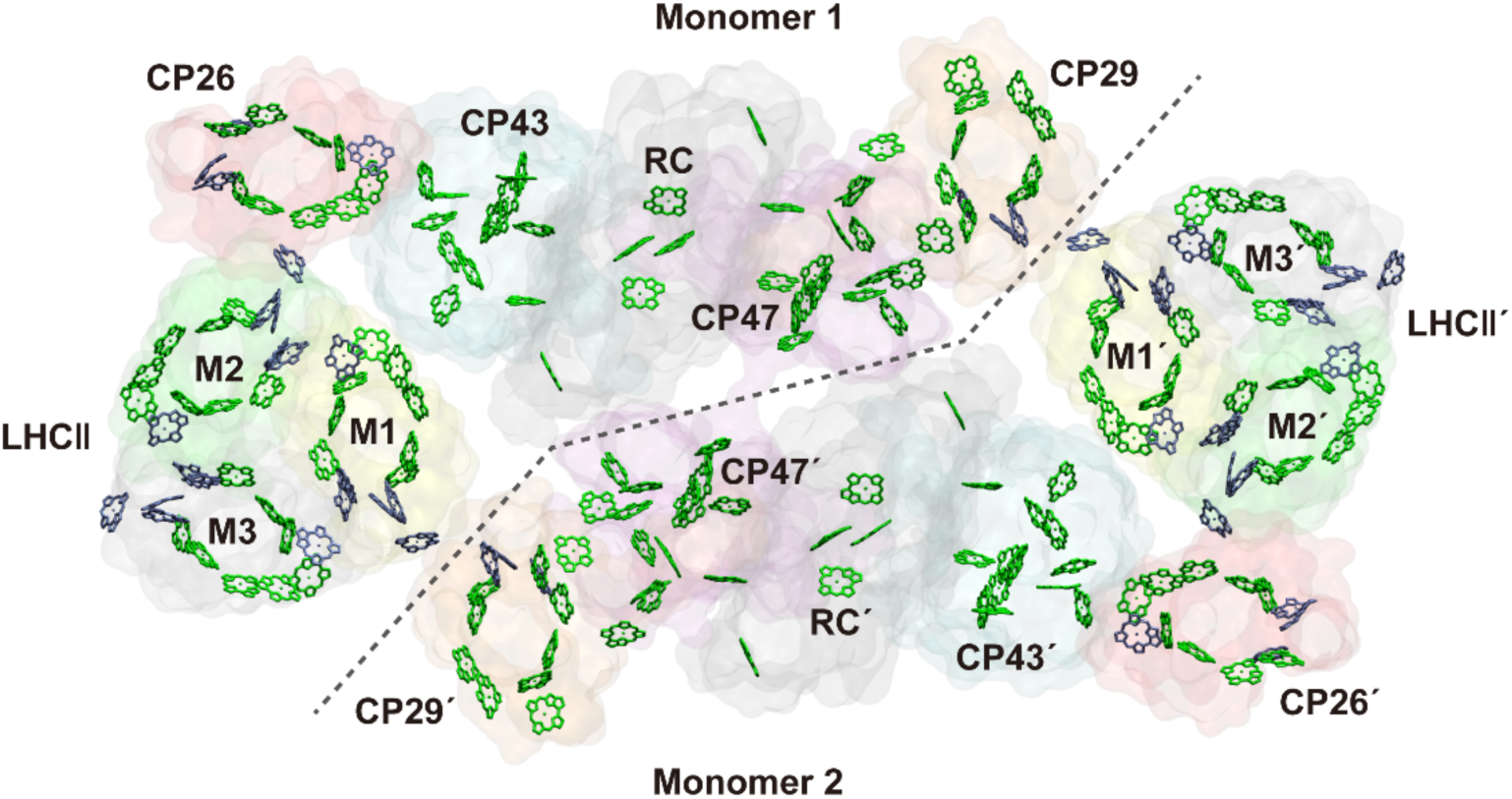
Schematic of protein subunits and pigment arrangement in the C2S2-type PSII-LHCII from spinach (PDB: 3JCU). Protein subunits are visualized using the QuickSurf drawing method with transparent materials. CP26 subunits are colored red, CP29 in orange, and LHCII monomers 1-3 in yellow, green, and gray, respectively. CP43 appears in blue, CP47 in purple, and the RC in dark gray. Pigment molecules are depicted via the licorice drawing method. Chlorophylls (Chls) a are shown in green, and Chls b in ice blue. A black dashed line marks the boundary between the two monomers.

Here, we use these simulations to explore the energy landscape and transfer pathways of PSII. By integrating MD, QM/MM calculations, and network analysis, we map EET within and between subunits, identify key pigments and dominant routes, and analyze how PSII achieves efficient, regulated energy flow despite a flat gradient. Our findings provide mechanistic insight into PSII’s EET strategy and may guide the design of artificial light-harvesting devices and crop improvement approaches.

## METHODS

### System setup and molecular dynamics simulation

The cryo-EM structure of a C_2_S_2_-type PSII-LHCII supercomplex from spinach resolved by Wei et al. (PDB ID: 3JCU^22^, resolution: 3.2 Å) was employed in this study. A 1 μs molecular dynamics simulation trajectory comprising approximately 1.2 million atoms, from our previous work^44^, was utilized for excited-state energy transfer calculations. As shown in Fig. 2A, we adopted a multiscale simulation framework. For conformational sampling, snapshots were extracted from the final 200 ns of the equilibrated trajectory at 10 ns intervals, yielding a total of 20 representative structures. Each snapshot was then subjected to 200 steps of molecular mechanics energy minimization to correct potential structural mismatches and prepare the system for subsequent QM/MM calculations.

**Fig. 2.**
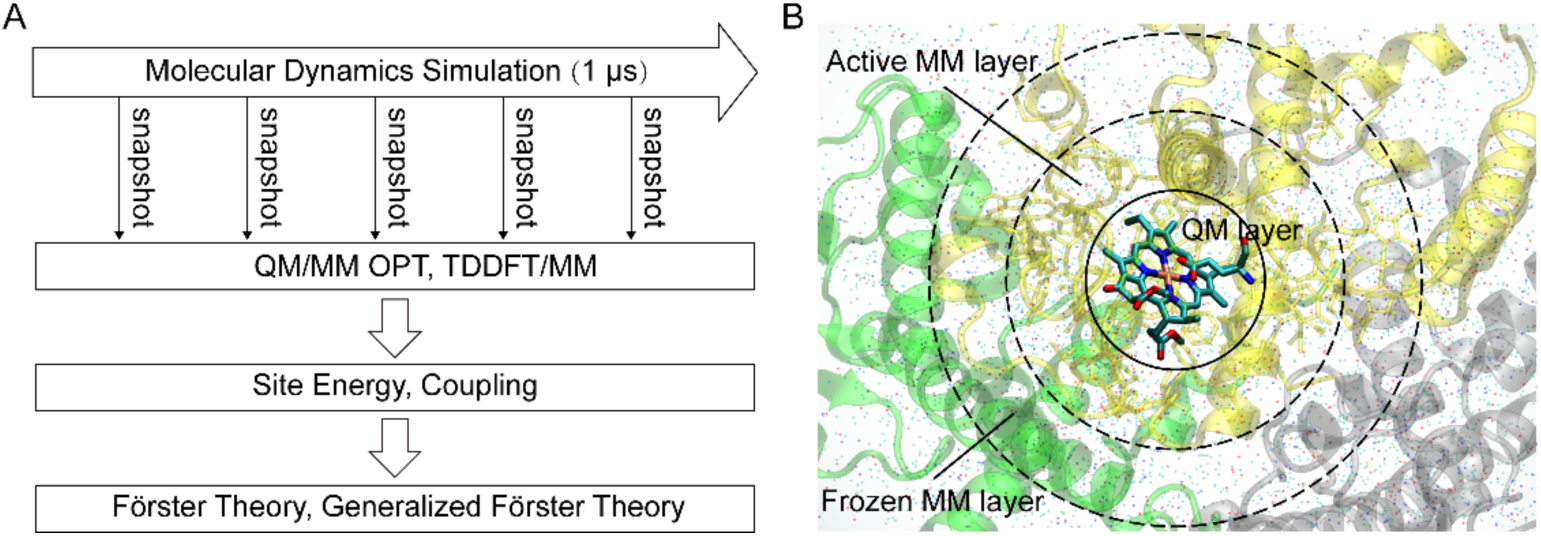
Schematic diagrams of multiscale simulation and three-layer QM/MM calculation. (A) The framework of multiscale simulation calculation. (B) the schematic diagram of the three-layer QM/MM calculation theme. The QM region, encompassing pigment molecules and their central ligands, is displayed in Licorice style. The active MM layer is rendered via the Licorice method with yellow transparent material, while the protein is visualized using the NewCartoon method with transparent material, maintaining color consistency with Fig. 1. Other components, including water molecules and cofactors, are presented in Point format.

### QM/MM setup and structure optimization

From each snapshot, a three-layer QM/MM calculation was performed for each pigment (Fig. 2B). In this model, the system was partitioned into three regions: the QM layer, the non-frozen MM layer (active MM layer), and the frozen MM layer. The QM layer consists of a chlorophyll molecule with the phytylchain cut at the first carbon-carbon bond after the ester group, and the magnesium coordinating residue^45^. These residues located within a 3 Å cut-off radius of the pigment molecule. The non-frozen MM region, treated using molecular mechanics force field, as described in our previous work^44^ encompassed all atoms within a 7 Å radius of the QM atoms and was allowed to relax during geometry optimization. The frozen MM region included all remaining atoms within 40 Å of the pigment molecule, including protein, solvent, and cofactors; these atoms were kept fixed throughout the QM/MM calculations to maintain the structural environment. Employing this approach, we systematically extracted a total of 4,120 QM/MM models from the 20 sampled representative structures for QM/MM optimization. For each of these models, a comprehensive consideration was given to the non-frozen and frozen MM regions, as well as the QM atoms. Such an elaborate procedure ensured that each structure for QM/MM optimization was configured to accurately represent the complex interactions within the system^30, 46^. Considering the trade-off between computational cost and accuracy, each QM/MM geometry optimization was conducted in two sequential steps. In the first step, the active layer was optimized for 400 steps using the same force field employed in the MD simulations. In the second step, the QM region was optimized using the semiempirical AM1 method^47^, with the root-mean-square (RMS) gradient convergence threshold set to 0.01 kJ/mol/Å. The final optimized structures were then used for subsequent excited-state calculations. All QM/MM calculations were performed using a locally modified version of the pDynamo2 program^48, 49^, with external coupling to the BDF program package^50^ for quantum chemical computations.

### Theoretical calculation of site energies and electronic couplings

The electronic properties of the PSII-LHCII supercomplex are described using the Frenkel exciton model. In this framework, the total electronic Hamiltonian of the system is approximated as sum over single-pigment excitations and the corresponding electronic couplings among them, as shown in Equation ^28, 51^:

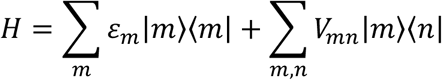

Here, the site energy *ε*_*m*_ corresponds to the excitation energy of pigment *m* in its first excited electronic state. The basis state |*m*⟩ represents the configuration in which all pigments remain in their electronic ground states, except for pigment *m*, which is in the excited state. The electronic coupling term *V_mn_* describes the interaction between excitonic states |*m*⟩ and |*n*⟩, arising from the Coulombic interaction between the transition densities of pigments *m* and *n*.

Gaussian16^52^ software was used to calculate the site energies, CAM-B3LYP/6-31G*^53^ level of theory was used for MM/TDDFT calculations. The electronic coupling term between pigments was calculated by the TrEsp (transition charge from electrostatic potential) method^54, 55^ and Multiwfn program^56^.

### Inter-pigment energy transfer rate and inter-subunit energy transfer rate calculations

Two theoretical methods were employed to calculate the energy transfer rates between pigments and between subunits, respectively. For the energy-transfer rate T (in ps^−1^) among pigment molecules, Förster theory^57, 58^ is utilized for calculations. In contrast, energy transfer between pigment clusters (i.e., subunits) was evaluated using generalized Förster theory^59, 60^. According to this theory, the energy transfer rate *k*_*DA*_ between a donor pigment cluster D and an acceptor pigment cluster A is computed based on their electronic coupling and spectral overlap. Detailed equations and computational scripts can be found in our previous work^45, 61, 62^.

## RESULTS AND DISCUSSION

### Energy landscape of site energies

Accurate determination of the site energy landscape is essential for understanding EET in photosynthetic complexes. The protein environment modulates pigment excitation through electrostatic interactions, structural distortions, coordinating residues, and hydrogen bonding^14, 63–66^. To capture these effects, we employed a QM/MM approach to calculate site energies for 206 pigments across 20 representative conformations. For comparison, excitation energies were also computed for the isolated QM regions without the surrounding protein environment. In total, 8,240 excited-state calculations were performed, and the final site energy of each pigment was obtained by averaging over the 20 conformations. Complete datasets are provided in Tables DS1-DS3 of the Extended Data.

To evaluate the influence of conformational dynamics, we analyzed the standard deviations (SDs) of site energies from both QM/MM and QM calculations (Table S1). For QM/MM, SDs ranged from ± 0.005 to ± 0.026 eV, with a mean of 0.014 eV; for QM, values ranged from ± 0.005 to ± 0.023 eV, with a mean of 0.013 eV. These narrow distributions indicate that conformational changes during dynamics exert only minor effects on pigment site energies, reflecting strong conformational constraints imposed by the protein scaffold. For comparison, Cardoso et al. reported a SD of 0.041 eV for B800 pigments, considerably larger than our values^14^. The reduced fluctuations observed in PSII thus further confirm the restraining role of the protein matrix on pigment conformations.

Fig. 3A compares the site energies obtained from QM/MM and QM calculations. The x-axis represents the average site energies of 206 pigments calculated using the QM/MM approach, while the y-axis shows the corresponding values from the QM-only calculations. The data points from both methods are closely aligned along the diagonal, indicating strong agreement between the two approaches. The typical variation is within ± 0.04 eV. Statistical analysis using Student’s t-test further confirms that the differences between the QM and QM/MM results are not statistically significant (p > 0.05).

**Fig. 3.**
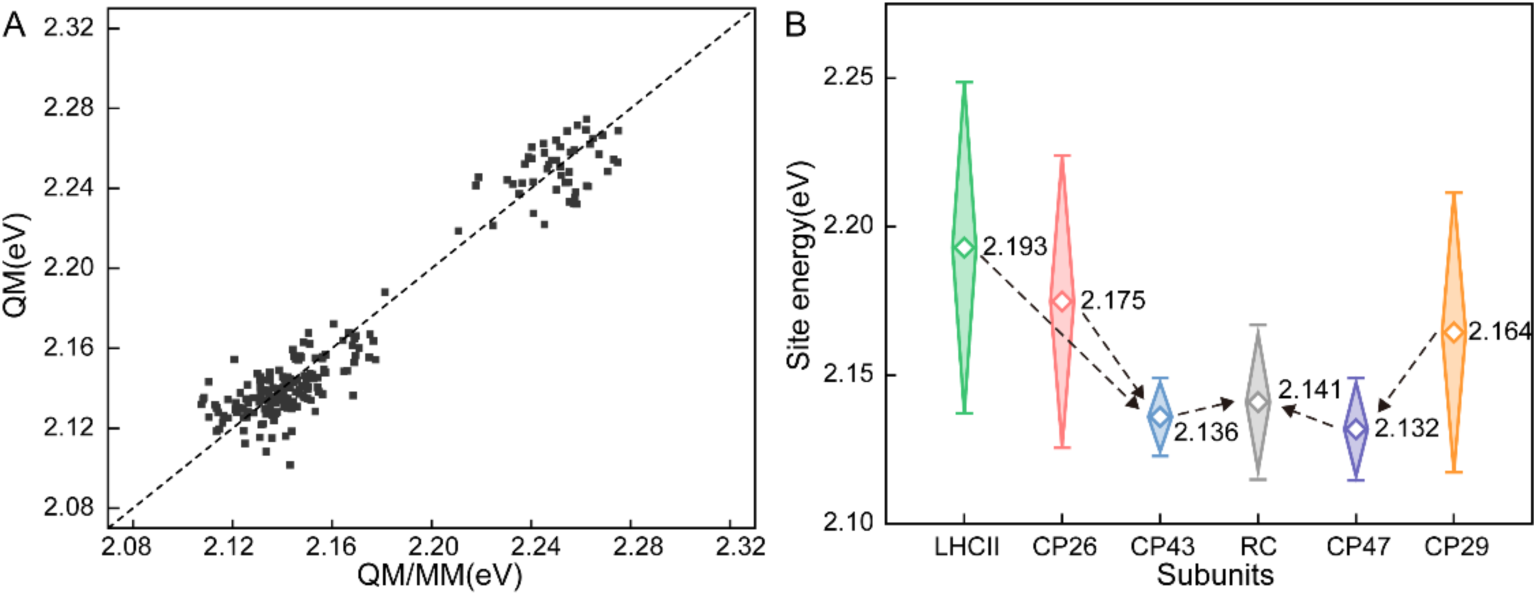
Analysis of site energies based on pigments and subunits. (A) Comparative analysis of site energies for 206 pigments in PSII-LHCII using QM/MM and QM methods. The x-axis shows average site energies of 206 pigments calculated by QM/MM methods, and the y-axis denotes average site energies of the same pigments by QM methods. The diagonal dashed line indicates sites where the QM/MM and QM calculations are in agreement. (B) Diamond box plots of site energies for in C2S2-type PSII-LHCII subunits derived from QM/MM method. Hollow diamonds indicate mean values, with boxes representing standard deviations (SD). Subunits are color-coded: CP26 (red), CP29 (orange), LHCII (green), CP43 (blue), CP47 (purple), and the reaction center (RC, dark gray). Black dashed arrows illustrate proposed energy transfer pathways from the peripheral antenna (LHCII/CP26/CP29) through the inner antenna (CP43/CP47) to the RC.

In fact, the primary difference between the QM and QM/MM methods arises from the electrostatic influence of the surrounding protein matrix, which has been well documented in previous studies^63, 67–69^. In our calculations, to isolate the impact of the electrostatic environment, we ensured that the QM and QM/MM methods utilized identical pigment conformations (i.e., the same molecular structure), thus eliminating conformational variation as a confounding factor. Therefore, we conclude that the direct electrostatic contribution of the protein matrix to the pigment site energies is relatively minor. Of course, this does not imply that the protein environment has no effect on the site energies of pigment molecules. Instead, the primary role of the protein environment lies in modulating the pigment conformations, which in turn affects their excitation properties. Accordingly, in the subsequent analyses, we use only the site energies obtained from the QM/MM calculations.

Fig. 3B illustrates diamond box plots (with diamond markers for means) of the site energy distribution across pigments, obtained by averaging the site energies within each subunit of the PSII-LHCII supercomplex. This analysis provides insights into the directionality of excitation energy transfer and underscores the functional specialization of individual subunits^37^. The average site energies exhibit a decreasing trend from the peripheral antenna to the core antenna, following the order LHCII/CP26 > CP43 and CP29 > CP47, which is consistent with the principle of downhill excitation energy transfer^37, 70–72^. However, the average site energies of CP43 and CP47 are slightly lower than that of the RC, suggesting an apparent uphill energy transfer. Nevertheless, the energy differences are relatively small, at only 0.009 eV and 0.005 eV, respectively. To ensure that these differences are not due to variations in pigment composition across subunits, we performed a control analysis considering only Chl a pigments (see Supporting Information, Table S2). In this case, the inter-subunit site energy gradient becomes significantly flatter but still retains the overall downhill trend described above.

We further analyzed the inter-subunit site energy gradient using QM-only calculations (Supporting Information, Fig. S1), which revealed the same trend as observed in the QM/MM calculations. This peculiarity was reported in 2016, based on a 12 Å resolution model, when Kreisbeck and colleagues suggested that an energy barrier must be overcome during transfer from PSII to the RC^37^. Our current findings are consistent with this observation, indicating that the presence of such a barrier is a recurrent feature of PSII energy transfer. However, as discussed in later sections, our analyses suggest that uphill transfer may not be essential.

### Energy transfer network within PSII Complex

The pairwise EET rates among 206 pigments were calculated using Förster theory to investigate dynamics at the pigment level, and a pigment transfer network was constructed for further analysis (Fig. 4). The overall distribution of EET rates in the PSII–LHCII system is shown in Fig. 4A. Most transfer events occur on the sub-picosecond to picosecond timescale, forming an extensive EET network. The fastest EET channels (> 1 ps^−1^) are concentrated in strongly coupled regions within subunits (Supporting Information, Fig. S2) and at some specific interfaces between peripheral and inner antenna pigments. For example, within an LHCII monomer, transfers among a611-a612 and a610 exceed 1 ps^−1^, due to the strongly-coupled within the subunit (Fig. S2 and Fig. 4A). Moreover, moderately coupled pigment pairs at the LHCII-CP43 interface enable efficient delivery of excitation from peripheral to core antennas (Fig. S2 and Fig. 4A).

**Fig. 4.**
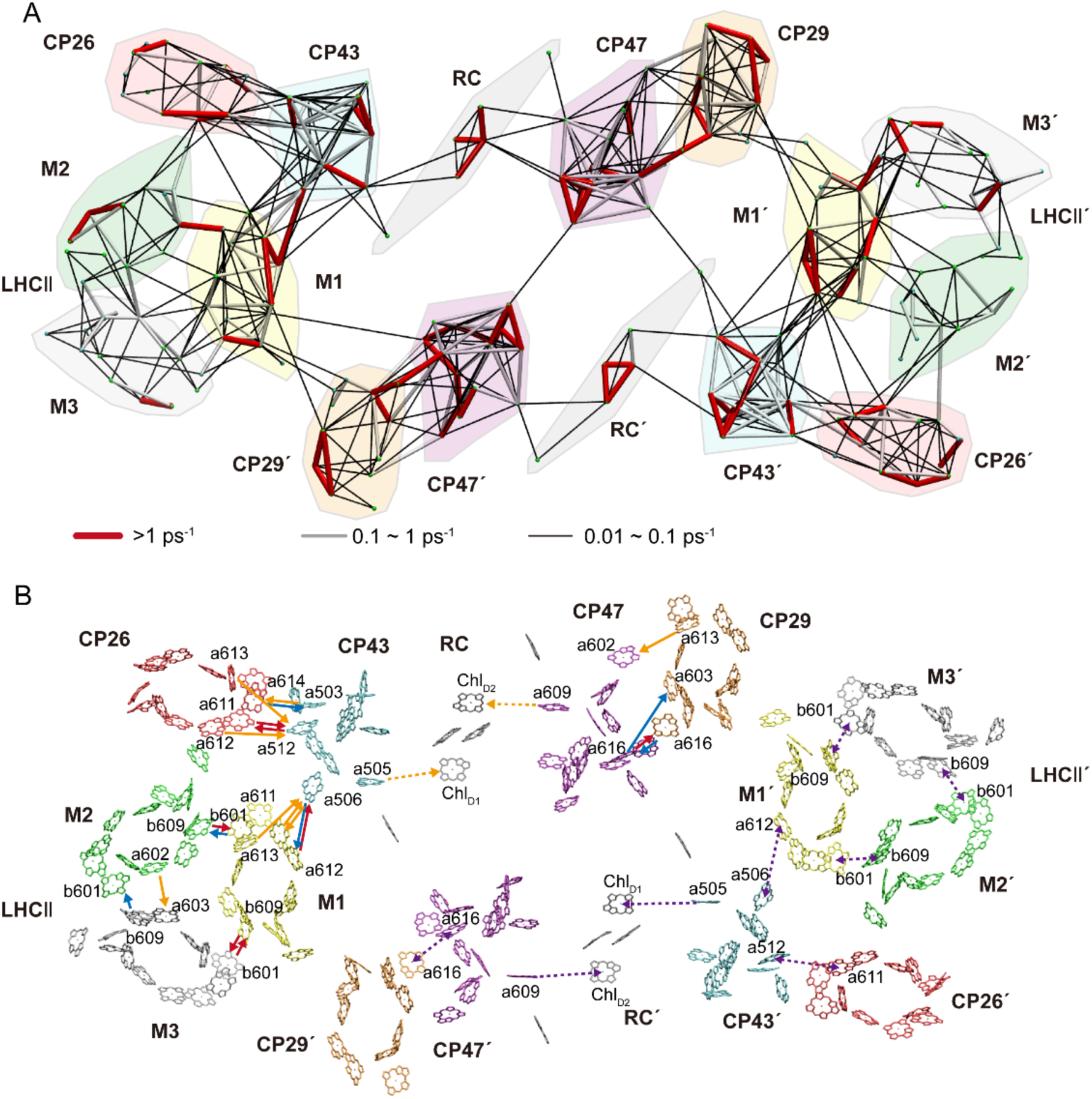
FRET network and excitation energy transfer (EET) pathways in PSII-LHCII supercomplex. (A) The map of inter-pigment EET rates. Inter-pigment rates slower than 0.01 ps^−1^ are omitted. (B) EET pathways between the pigments are visualized from the stromal side, with time constants (τ, ps) indicated by arrows. EET pathways from the peripheral antenna to the inner antenna are indicated by solid arrows categorized by time constants: red arrows denote τ < 2 ps, blue arrows represent 2 ≤ τ < 5 ps, and orange arrows signify 5 ≤ τ ≤ 10 ps. Inner antenna to RC transfers are shown as orange dashed arrows. The fastest EET pathways between subunits are indicated by a purple dotted arrow in monomer 2. The displayed time constants (τ, ps) represent the inverse of the energy transfer rates (τ=1/***T_m→n_***, in ps).

Intermediate EET processes (0.1-1 ps^−1^) are distributed both within and between subunits, but form a relatively sparse network. Within subunits, these channels connect clusters involved in the fast pathways, while between subunits they provide complementary routes that maintain basic connectivity. Slow transfers (0.01-0.1 ps^−1^) further extend the network by linking pigment pairs not involved in faster processes, particularly establishing connections between the inner antenna and the RC. Together, these multi-timescale pathways construct a complete EET network that ensures smooth energy delivery to the RC, where charge separation is initiated^73–75^.

Analysis of inter-subunit transfer pathways reveals that multiple parallel routes exist across subunit interfaces (Fig. 4B). For example, between CP26 and CP43, three parallel transfer paths were identified: a611 (a612) →a512, a613→a512, and a614→ a503, with time constants of < 2 ps, < 10 ps, and < 5 ps, respectively. A similar pattern is observed at the interface between LHCII and CP43, where multiple fast parallel transfer routes are also present. The presence of several rapid pathways at these interfaces likely enhances the robustness and efficiency of excitation energy delivery, ensuring that energy transfer can proceed efficiently even if one route is transiently hindered or energetically unfavorable. In previous studies based on pigment distances or theoretical calculations with empirical parameters, research on the resolved structures of PSII in different species has indicated that the energy transfer between the peripheral antenna and the inner antenna involves multiple possible pathways^21–23, 27, 76^. Notably, although multiple pathways exist for inter-subunit transfer, most converge on one or two key pigments located at the interfaces, such as a506 at the LHCII-CP43 interface and a512 at the CP26-CP43 interface. These interface pigments likely function as energy-funneling sites, acting as functional hotspots that integrate excitation energy from multiple incoming routes and channel it toward the reaction center. Their strategic positioning may also facilitate dynamic regulation of inter-subunit energy flow, allowing excess excitation to be redirected under high-light conditions as part of photoprotective mechanisms^41^.

To investigate the relationship between the site energy landscape and the energy transfer pathways, the excitation energy transfer network of the CP43-RC-CP47 core complex was analyzed using a spatial representation, in which the x and y coordinates reflect the relative positions of pigments and the z coordinate represents their site energies (Fig. 5). This approach provides a comprehensive view of the spatial distribution of PSII core pigments and the relative differences in their site energies. For transfers between the inner antennas and the RC, only pathways with time constants shorter than 40 ps were included. For intra-subunit connections, the time constants were restricted to less than 20 ps.

**Fig. 5.**
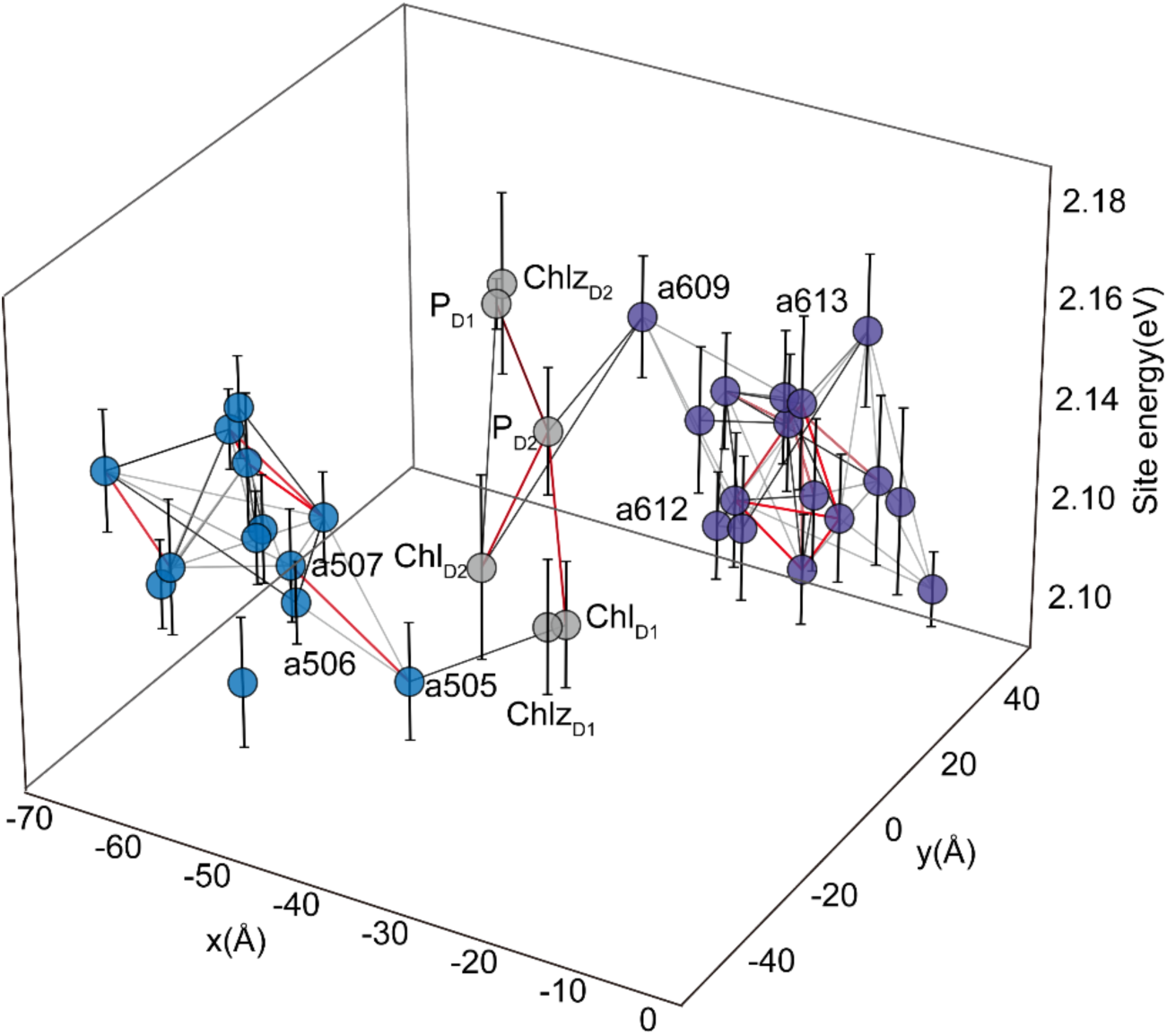
Pigment site energies and FRET network in the CP43-RC-CP47 Core Complex. Pigment positions are represented by x and y coordinates of magnesium atoms with circular markers: blue circles for CP43 pigments, purple circles for CP47 pigments, and gray circles for RC pigments. The z-axis denotes the average site energy of each pigment, with error bars indicating the standard deviation of site energies. Energy transfer time constant ranges are visualized with color-coded lines: red for τ ≤ 1 ps, gray for 1 < τ ≤ 10 ps, and black for 10 < τ ≤ 20 ps pathways. Energy transfer from the inner antenna to the RC is indicated by black lines, with data restricted to τ < 40 ps.

As shown in Fig. 5, using the cutoff, the pigments of PSII core form a complete energy transfer network. Based on the collaborative analysis of site energies and network structures, this difference stems from distinct rate-limiting mechanisms on the two sides. On the D1 side of the photosynthetic RC, a relatively efficient energy transfer process unfolds. Energy initially enters CP43 and converges to a505 through a series of consecutive downhill transfers. This is a crucial step as it effectively channels the energy towards a focal point within CP43. Subsequently, the energy is transferred to the Chl_D1_ in the RC within a spatial distance of 27.64 Å. This specific energy transfer route from CP43 to Chl_D1_ on the D1 side is not only a matter of efficient energy conveyance but also holds significant functional implications. It forms the basis for the primary reaction branch in the photosynthetic process. The fact that energy converges on a505 in CP43 and then moves on to Chl_D1_ primes the system for the subsequent electron transfer events^69, 77, 78^. As previous studies have indicated, the pigment Chl_D1_ on the active branch has the lowest site energy within PSII^13, 77, 79^. This low site energy endows Chl_D1_ with the ability to serve as the primary electron donor. In essence, the energy transferred to Chl_D1_ via the a505 pathway enables Chl_D1_ to initiate the electron transfer chain, which is fundamental for the conversion of light energy into chemical energy. In contrast, on the D2 side, CP47 has to overcome an “uphill” energy gradient to transfer energy to a609. Subsequently, following the downhill energy transfer pattern, it transfers the energy to Chl_D2_ of D2, with a distance of 26.4 Å between them. This differential energy transfer between the D1 and D2 sides of PSII is a crucial aspect of its functionality. The D1 side, characterized by a downhill energy transfer pathway, enables energy delivery to the RC. This efficient transfer primes the RC for swift initiation of primary photochemical reactions. Conversely, the “uphill” energy gradient on the D2 side, while seemingly a hindrance, plays an essential role in the overall energy management of PSII. It acts as a regulatory mechanism, controlling the rate at which energy reaches the RC. By preventing an excessive influx of energy, it safeguards the RC from over-excitation, which could otherwise lead to the generation of reactive oxygen species and subsequent photodamage^43, 80, 81^. In summary, the existence of these two differential energy transfer pathways within PSII represents an evolutionary adaptation that optimizes light-harvesting efficiency while maintaining the stability and integrity of the photosynthetic apparatus. This intricate balance between efficient energy transfer on the D1 side and regulated energy input on the D2 side is fundamental to the overall success of photosynthesis under diverse environmental conditions.

To test this inference, we examined pigments a609, a612, and a613 and plotted Gaussian distributions based on site energy data from 20 dynamic conformations (Fig. 6A). The results show substantial overlap among these distributions: a612 is mainly distributed between 650-700 nm, a609 between 640-680 nm, and a613 spans the widest range, 640-690 nm. This overlap indicates intersecting electronic states. Notably, the broad distribution of a613 reflects its strong structural fluctuations, which can transiently raise its site energy above that of a609, thereby providing a thermodynamic driving force for energy transfer to a609. By analyzing site energy time series from the last 200 ns of MD simulations (Fig. 6A), we further confirmed the dynamic nature of site energy fluctuations. Sites a609, a612, and a613 all exhibited irregular variations: a609 decreased markedly at 30, 60, 160, and 190 ns; a612 was usually lower than a609 but exceeded it at 30 and 190 ns; and a613 showed the strongest fluctuations, surpassing a609 at multiple points (30, 50, 60, 130, 160, 190 ns). From the coupling matrix (Fig. S2, Extended Data Tables DS4), a612 and a613 are moderately coupled to a609 (11-13 cm^−1^). Thus, whenever a612 or a613 attains higher site energy than a609, a transient downhill gradient emerges, enabling efficient energy transfer to a609.

**Fig. 6.**
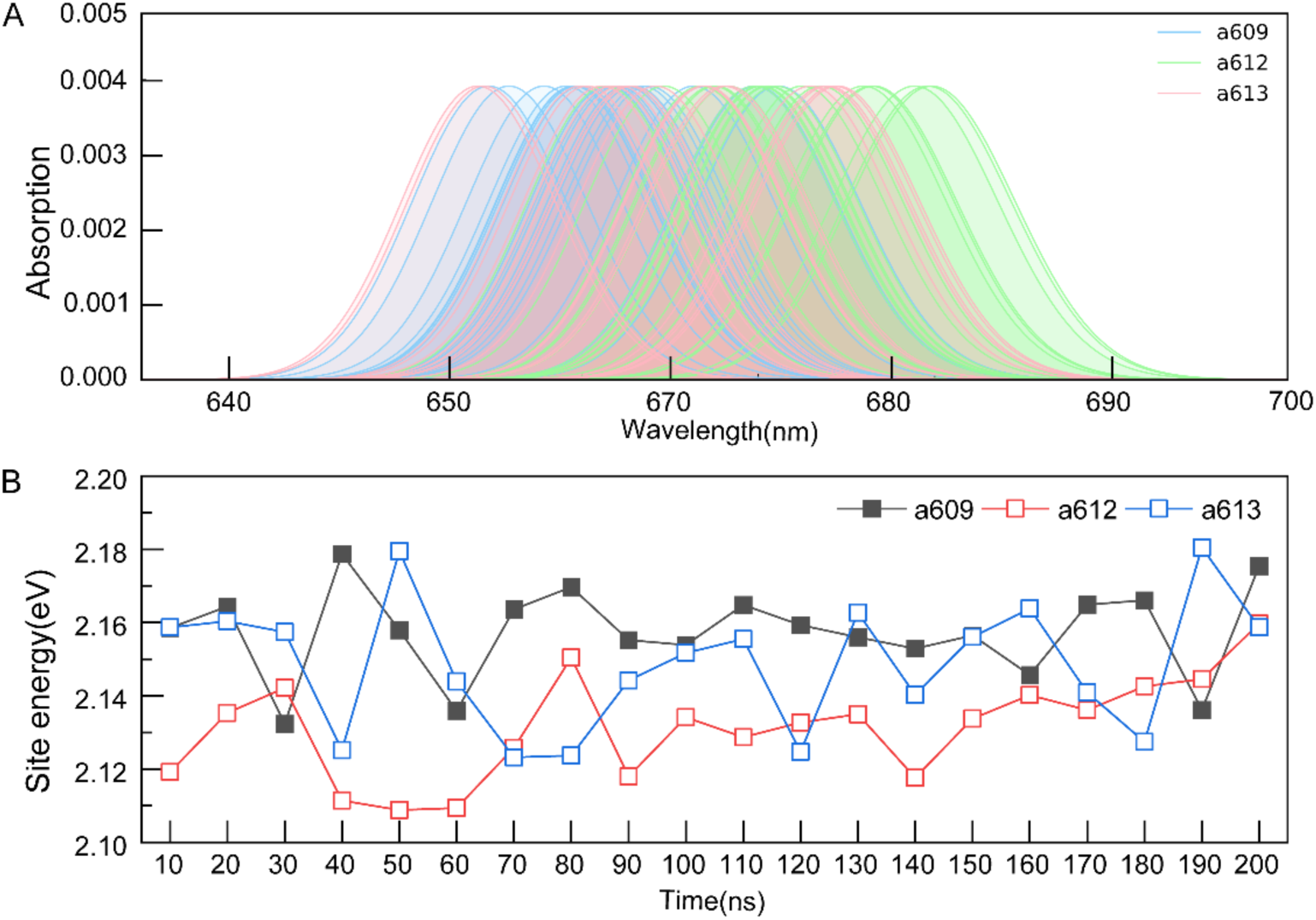
Gaussian spectra and temporal fluctuations of site energies of a609 a612 and a613 pigments in CP47. (A) Gaussian spectra of a609 a612 and a613 pigments in CP47. The spectrum of each pigment is plotted based on the site energies calculated from its 20 conformations, with a609 spectrum in light blue a612 in light green and a613 in light pink; all spectra have been applied with a 0.285 eV shift correction. (B) Temporal fluctuations of site energies for a609 a612 and a613 pigments within CP47. The data are derived from the last 200 ns of the simulation trajectory with a sampling interval of 10 ns per frame.

In summary, the apparent uphill transfer from CP47 to the RC arises from dynamic downhill processes driven by structural fluctuations. Although the average site energy of a609 is higher than that of other CP47 pigments, a strongly-coupled pigment cluster in CP47 can efficiently gather and transfer energy to a612 and a613. Structural fluctuations dynamically reshape relative site energies, and at multiple time points a612 and a613 surpass a609, allowing downhill transfer. This mechanism demonstrates that CP47 still follows the overall downhill transfer principle: the uphill phenomenon is a static artifact of averaged site energies, while dynamic fluctuations ensure the thermodynamic feasibility of energy flow.

### Inter-Subunit Energy Transfer Network

Using generalized Förster theory, we computed EET time constants between PSII subunits to characterize the efficiency and hierarchy of transfer pathways to the RC (Fig. 7). Only pathways with time constants shorter than 250 ps are considered, consistent with a previous study^37^. Three parallel transfer routes were identified. Path 1 (D1 side) proceeds from CP26 to CP43 and then to the RC (128 ps), the fastest among them. Path 2 (D2 side) extends from CP29 to CP47 to the RC (229 ps). Path 3 begins in LHCII, passes through CP43, and reaches the RC (141 ps). All fall within the functional timescale reported by Kreisbeck^37^. Earlier studies on peas and green algae, based on empirical parameters, also identified three major routes but reported CP29→CP47 as the fastest^31^. In contrast, our analysis indicates that the CP26 pathway is slightly faster, likely reflecting the improved accuracy of our structural and theoretical framework.

**Fig. 7.**
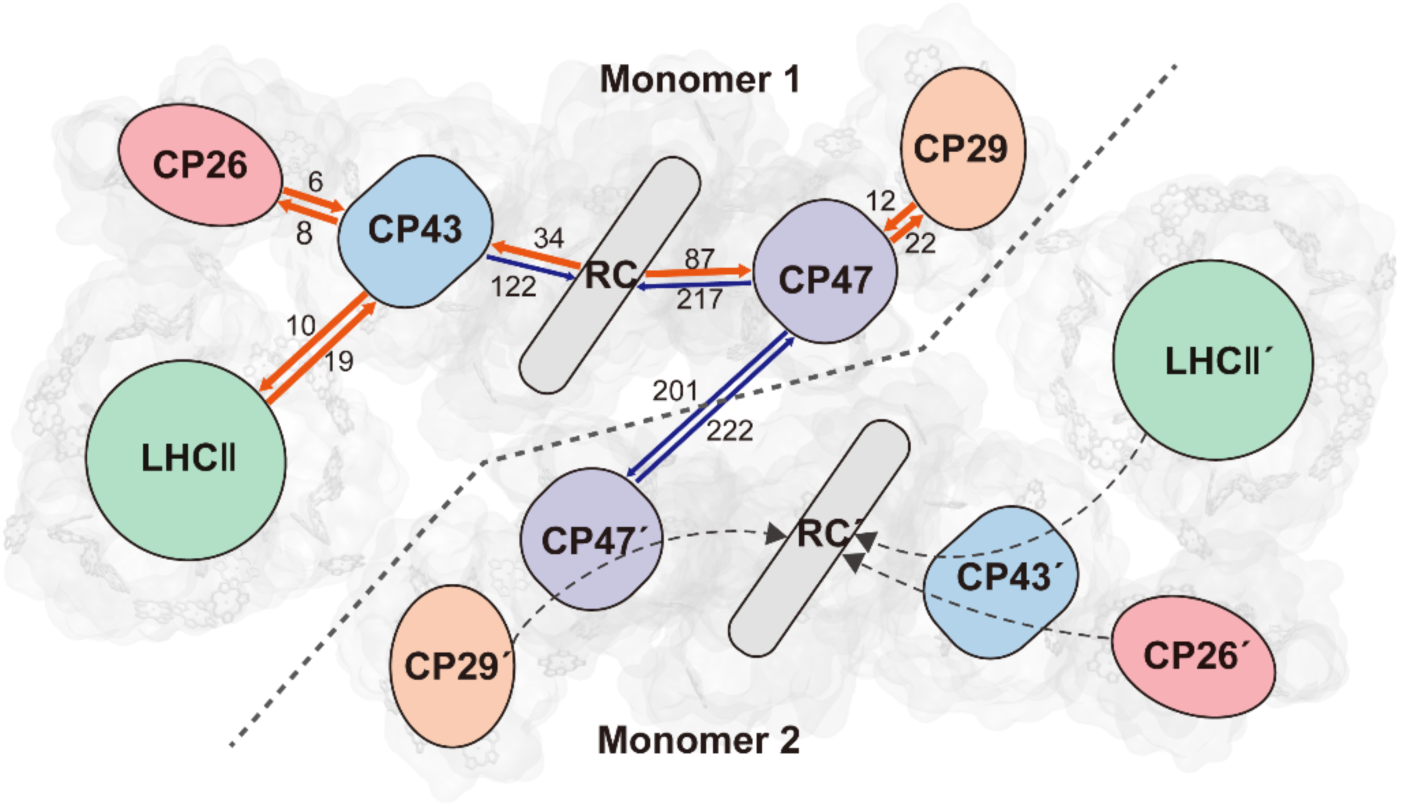
Schematic diagram illustrating energy transfer time constants and pathways between subunits calculated using generalized Förster theory. Time constants (τ, in picoseconds, ps) for energy transfer between subunits are labeled on the Monomer 1 side: orange arrows denote pathways with τ ≤ 100 ps, and blue arrows denote pathways with τ ≤ 250 ps. Major excitation energy transfer pathways from the light-harvesting complexes to the reaction center (RC) on the Monomer 2 side are indicated by gray dashed arrows. The displayed time constants (τ, ps) represent the inverse of the energy transfer rates (***k_DA_*** = 1/τ, in ps^−1^).

All three routes exhibit rapid bidirectional transfer between peripheral and core antennas. For CP26–CP43, forward transfer occurs in 6 ps and reverse in 8 ps, consistent with Yang et al.^43^. CP29-CP47 shows 12 ps forward and 22 ps reverse transfer. For LHCII-CP43, forward transfer requires 19 ps, while reverse is faster at 10 ps, shorter than values reported previously^29,30^. Such bidirectional dynamics likely contribute to photoprotection by allowing excess excitation to be redirected from the core back to peripheral antennas for dissipation^41, 82, 83^.

## CONCLUSIONS

In this work, we combined QM/MM calculations, Förster theory, and network analysis to investigate EET in the PSII-LHCII supercomplex. Our site energy calculations for 206 pigments showed strong consistency between QM/MM and QM-only methods, with typical variations within ± 0.04 eV. The protein matrix contributes only weak direct electrostatic effects but plays a dominant role in constraining pigment conformations, which in turn modulate excitation properties. The overall site energy gradient decreases from peripheral to inner antennas and toward the RC, although CP43 and CP47 appear slightly lower than the RC. This apparent uphill feature, reported previously, is largely a static artifact, as dynamic structural fluctuations generate transient downhill gradients that ensure thermodynamic feasibility.

Förster-based calculations established a dense multi-timescale EET network spanning 206 pigments. Fast transfers (< 1 ps^−1^) dominate within subunits and at antenna interfaces, while slower channels maintain overall network connectivity. At several interfaces, such as CP26-CP43 and LHCII-CP43, multiple parallel routes converge on a few key pigments that serve as energy-funneling hotspots. This structural organization enhances robustness and provides flexibility, allowing excitation to be redirected when individual routes are hindered.

Analysis of the PSII core revealed distinct strategies on the D1 and D2 sides. On the D1 side, energy flows smoothly downhill through CP43 and converges at a505 before being delivered to Chl_D1_, the primary electron donor. On the D2 side, CP47 shows an apparent uphill step before transfer to Chl_D2_. However, structural fluctuations and moderate pigment couplings dynamically reshape relative site energies, enabling effective downhill transfer to a609 and then to Chl_D2_.

Generalized Förster theory identified three major transfer pathways to the RC: CP26→CP43→RC (128 ps), CP29→CP47→RC (229 ps), and LHCII→CP43→RC (141 ps). All fall within the functional timescale of PSII. Importantly, all three pathways exhibit bidirectional transfer, with reverse flows often faster than forward ones—for instance, RC→CP43 occurs in 34 ps compared to 122 ps for CP43→RC. Such rapid reverse transfers provide a key photoprotective mechanism, allowing surplus excitation to be shunted back toward the antennas for dissipation.

Overall, our findings demonstrate that the PSII-LHCII supercomplex integrates structural constraints, redundant pathways, and dynamic regulation to achieve both efficient energy delivery to the RC and robust photoprotection. This dual strategy ensures that photosynthesis maintains efficiency and stability under diverse and fluctuating light environments.

## Author Contributions

JG: idea conceptualization. JG, LHB and LNL: project administration and funding acquisition. JPG, FX, RCM and JG: methodology, data analysis, and validation. JPG, JG, and LHB: manuscript writing.

## Notes

The authors declare no competing financial interests.

## ACKNOWLEDGEMENTS

This work was supported by the National Natural Science Foundation of China (21873034), Fundamental Research Funds for the Central Universities (2662024XXPY003 and 2662023XXPY006), Biotechnology and Biological Sciences Research Council (BB/W001012/1, BB/V009729/1, BB/Y01135X/1, BB/R003890/1), Royal Society (URF\R\180030), and Leverhulme Trust (RPG-2021-286). The numerical computations were performed in Hefei Advanced Computing Center.

## AUTHOR INFORMATION

### Corresponding Authors

**Jun Gao** − Hubei Key Laboratory of Agricultural Bioinformatics, College of Informatics, Huazhong Agricultural University, Wuhan 430070, Hubei, China.

### Authors

**Lu-Ning Liu** − Institute of Systems, Molecular and Integrative Biology, University of Liverpool, Liverpool L69 7ZB, UK; Hubei Key Laboratory of Agricultural Bioinformatics, College of Informatics, Huazhong Agricultural University, Wuhan 430070, Hubei, China.

**Li-Hua Bie** − Hubei Key Laboratory of Agricultural Bioinformatics, College of Informatics, Huazhong Agricultural University, Wuhan 430070, Hubei, China.

**Jian-Ping Guo** − Hubei Key Laboratory of Agricultural Bioinformatics, College of Informatics, Huazhong Agricultural University, Wuhan 430070, Hubei, China.

**Fan Xu** − Hubei Key Laboratory of Agricultural Bioinformatics, College of Informatics, Huazhong Agricultural University, Wuhan 430070, Hubei, China.

**Rui-Chao Mao** − College of Chemical and Biological Engineering, Zhejiang University, Hangzhou 310058, China; Hubei Key Laboratory of Agricultural Bioinformatics, College of Informatics, Huazhong Agricultural University, Wuhan 430070, Hubei, China.

