## Supplemental file for "Multi-Scale Simulations of Excited-State Energy Transfer Pathways in the C2S2-Type PSII-LHCII Supercomplex of Spinach"

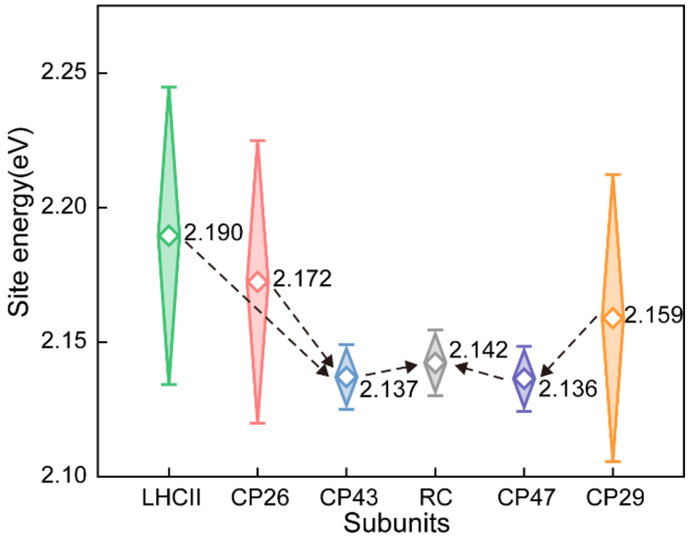


**Fig. S1 Diamond box plots of site energies for in $C_2S_2$-type PSII-LHCII subunits derived from QM method.** Hollow diamonds indicate mean values, with boxes representing standard deviations (SD). Subunits are color-coded: CP26 (red), CP29 (orange), LHCII (green), CP43 (blue), CP47 (purple), and the reaction center (RC, dark gray). Black dashed arrows illustrate proposed energy transfer pathways from the peripheral antenna (LHCII/CP24/CP29) through the inner antenna (CP43/CP47) to the RC.

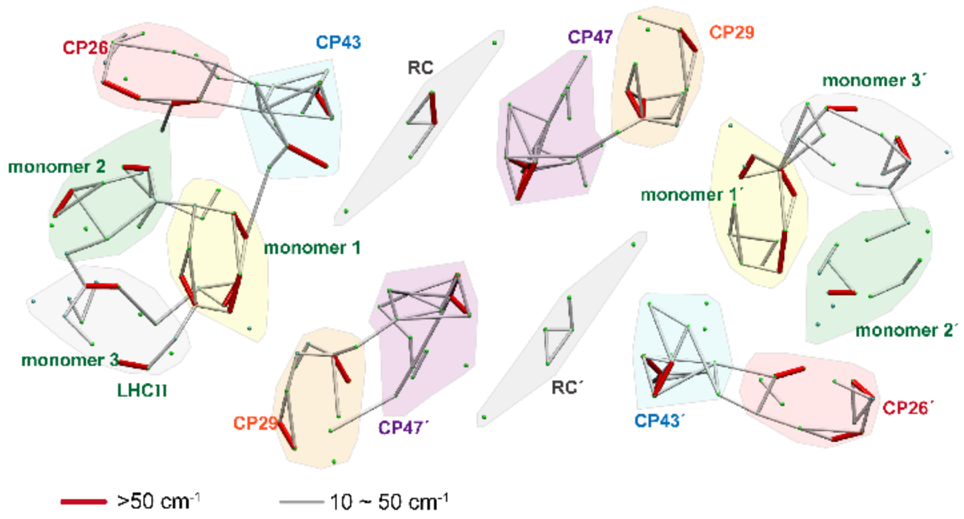


**Fig. S2 Schematic diagram of the pigment coupling network in PSII-LHCII supercomplex.** Inter-pigment couplings weaker than 10 $cm^{-1}$ are omitted.

**Tabel S1 Statistics of standard deviations of site energies for QM/MM and QM calculations on 206 pigment sites**

| | QM/MM | QM |
|---|---|---|
| Mean of Standard Deviation | 0.014 | 0.013 |
| Maximum of Standard Deviation | 0.026 | 0.023 |
| Minimum of Standard Deviation | 0.005 | 0.005 |

**Tabel S2 Statistical results of site energy of chl a in each subunit of $C_2S_2$-type PSII-LHCII**

| Protein | Average site energy(eV) | SD |
|---|---|---|
| LHCII trimer | 2.147 | 0.0136 |
| CP29 | 2.140 | 0.0080 |
| CP26 | 2.145 | 0.0171 |
| CP47 | 2.132 | 0.0158 |
| CP43 | 2.136 | 0.0117 |
| RC | 2.141 | 0.0231 |
